# Insulin-like Growth Factot-1 Supplementation Promotes Kidney Development and Alleviate Renal Inflammation in Preterm Pigs

**DOI:** 10.1101/2023.06.01.543191

**Authors:** Jingren Zhong, Thomas Thymann, Per Torp Sangild, Duc Ninh Nguyen, Tik Muk

**Affiliations:** Section for Comparative Paediatrics and Nutrition, Department of Veterinary and Animal Sciences, University of Copenhagen, Frederiksberg, Denmark; Department of Pediatrics, Odense University Hospital, Odense, Denmark; Department of Neonatology, Rigshospitalet, Copenhagen, Denmark

## Abstract

**Background:** Preterm birth and its associated complications cause disruption of normal prenatal renal development, leading to postnatal kidney injury and failure. Preterm infants are deficient in insulin-like growth factor 1 (IGF-1), a critical growth factor that stimulates tissue perfusion and development. Using necrotizing enterocolitis-sensitive preterm pigs as a model for preterm infants, we investigated whether IGF-1 supplementation during early life could improve kidney development and health.

**Methods:** Caesarean-delivered preterm pigs were allocated into two groups, either consistently receiving vehicle or IGF-1 immediately after birth for 5, 9 or 19 days. Postnatal age-matched term pigs were selected and served as term control on postnatal day (PND) 5, 9, and 19. Blood, urine and kidney tissue were collected for biochemical, histological and gene expression analyses.

**Results:** Preterm pigs showed impaired kidney development and increased kidney insults, as indicated by reduced average glomerular area, increased abnormal glomeruli percentage and increased markers of renal injury and inflammation compared to term pigs. IGF-1 supplementation significantly reduced the abnormal glomeruli percentage, renal injury and inflammation related markers, and up-regulated certain maturation-related genes on PND5.

**Conclusion:** IGF-1 supplementation supports kidney maturation and restoration of kidney insults after preterm birth in the early life of newborns.

**Impact:**

1. Preterm birth disrupts kidney development in preterm pigs.
2. Preterm birth leads to kidney injury and inflammation in preterm pigs.
3. IGF-1 supplementation might promote kidney maturation and alleviate preterm birth associated kidney injury and inflammation in preterm pigs.

## Introduction

Kidney develops in parallel with other organs (e.g. lungs, vascular tree) by branching morphogenesis that continues into the postnatal period ^1,2^. Over 60% of nephrons are formed during the last trimester of gestation ^3^ and the formation of new nephrons continues after birth ^4,5^. Preterm birth, which induces a series of whole-body complications ^6,7^, alters fetal renal development and forces the kidney to quickly adapt to the new extra-uterine environment. This may result in compromised nephrogenesis, renal insufficiency and ultimately renal failure ^1,8,9^. Preterm infants exhibit abnormal renal morphology ^10^, and insulted glomerular and tubular functions ^11-13^, predisposing them to neonatal acute kidney injury (AKI) in postnatal life (14). Yet, the underlying mechanisms and pathology of impaired renal development following preterm birth remain unclear ^14^.

The maturation of the kidney involves various processes, including branching of the ureteric bud and mesenchymal-epithelial transformation, which are regulated by a set of molecules and signaling pathways such as vascular endothelial growth factor A (VEGFA) and the glial-cell-line-derived neurotrophic factor (GDNF)/Ret proto-oncogene (RET) signaling pathway ^15-17^. Limited knowledge exists regarding how preterm birth and its associated complications may affect the renal development related signaling pathways. As nephrogenesis is sensitive to inflammatory insults ^18,19^, it is possible that impaired renal development is not only due to the immaturity but also the inflammation-related complications after preterm birth. Necrotizing enterocolitis (NEC) is an inflammatory disorder of the immature gut and may progress from mild mucosal lesions to severe mucosal destruction with systemic inflammation, sepsis and high mortality^20-22^. The latter severe complications with systemic inflammation could affect distant organs, such as the kidney, lung and brain, and may consequently contribute to AKI ^21,23,24^. Mild forms of neonatal gut complications (indicated by feeding intolerance) are widely present in preterm infants, but it is unclear if such mild local and/or systemic inflammatory effects may further predispose the population to AKI.

Insulin-like growth factor 1 (IGF-1) is a critical growth factor with a range of mitogenic, differentiating, and metabolic effects throughout the body in both fetal and postnatal life ^25^. IGF-1 is expressed in all fetal tissues and exhibits autocrine, endocrine, and paracrine actions mediated via both IGF-1 and insulin receptors ^26^. Around 80% of IGF-1 in circulation is bound to IGF-binding protein 3 (IGFBP-3) with an acid-labile subunit, which is capable of regulating IGF-1 functions in peripheral tissues ^27^. In preterm infants, circulating IGF-1 levels are considerably lower than their counterparts in utero, which may contribute to developmental deficiencies after birth, including kidney immaturity ^28-30^. Studies have demonstrated that exogenous IGF-1 administration may increase kidney weight in fetuses ^31^ and stimulate renal filtration and reabsorption ^29^.

The investigation of preterm birth associated kidney development requires appropriate animal models. Previous studies on baboons and lambs mainly focused on histological and physiological aspects of kidney maturation after preterm birth, without considering subsequent postnatal complications and further analysis on molecular changes ^9,14,19^. In this study, we used preterm pigs delivered at 90% gestation as a clinically relevant animal model of preterm infants to examine kidney development and insults specifically related to preterm birth-associated gut complications. Compared to previous studies focusing on mainly histological changes, this study placed greater emphasis on molecular changes in kidney tissue. We hypothesized that preterm birth is associated with delayed kidney maturation and increased inflammatory status in pigs, and that postnatal supplementation with IGF-1 would promote kidney maturation and mitigate the impact of preterm birth on kidney impairment.

## Methods

### Animal model and NEC evaluation

Animal studies were conducted in accordance with the European Communities Council Directive 2010/63/EU for the protection of animals used for experimental purposes and approved by the Danish Animal Experiments Inspectorate. The study design is briefly shown in Figure 1A, including three groups: term reference pigs, preterm control group, and preterm IGF-1 group.

**Figure 1.**
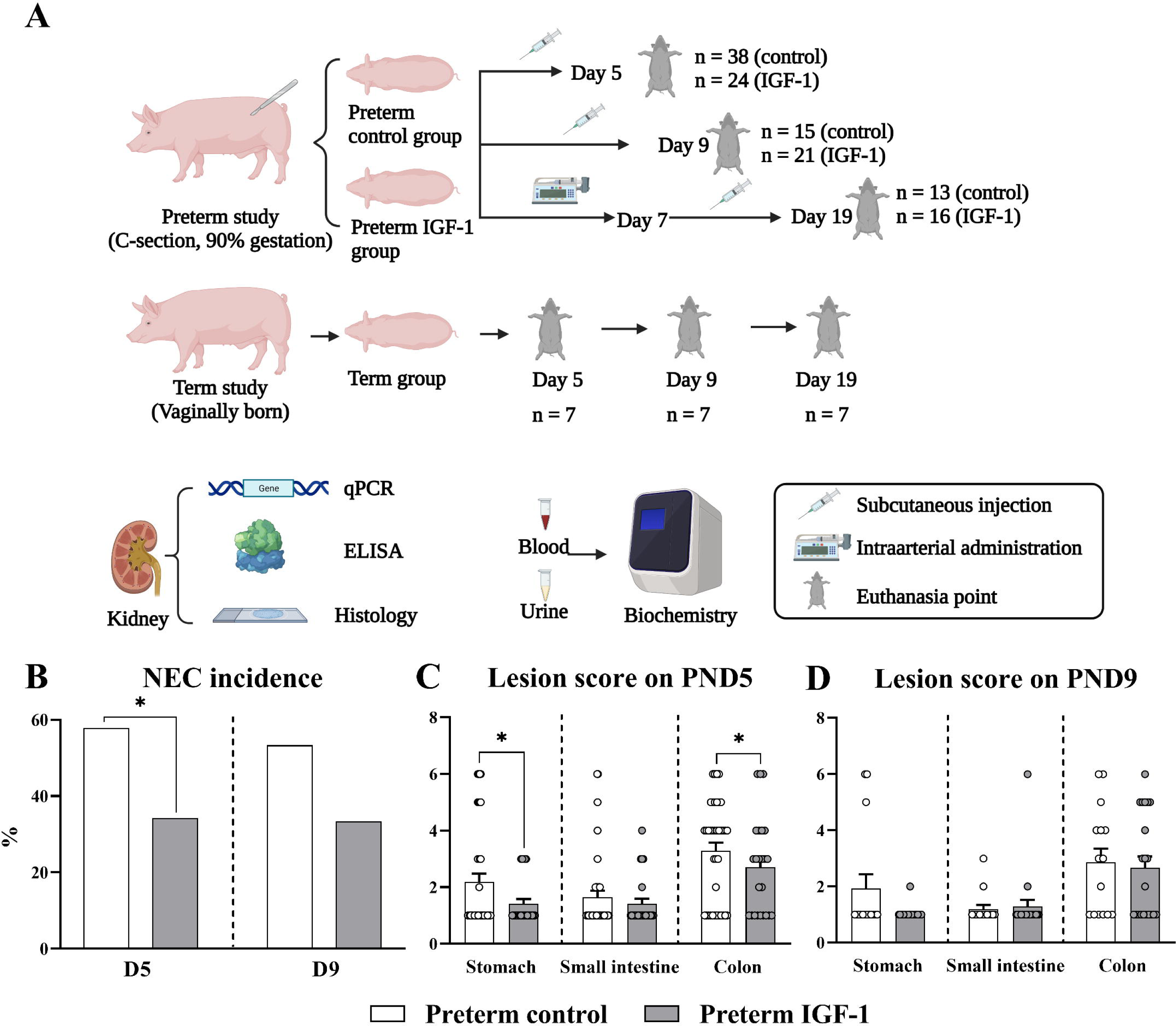
Study overview, Necrotizing Enterocolitis (NEC) conditions in preterm pigs. (A) Across three experiments, preterm pigs were treated with vehicle or rhIGF-1/IGFBP-3 for 5, 9 and 19 days. Vaginally born, farm-reared term pigs were selected at the matched postnatal age as the term reference. At the end of study, kidney, blood and urine samples were collected for further analysis. (B-D) NEC incidence and lesion distribution in preterm pigs. (B) NEC incidence in preterm and IGF-1 group on PND5 and 9 (NEC defined as a score of ≥4 in at least one region). (C and D) Individual lesion scores in stomach, small intestine and colon regions (highest recorded score in each region) on PND5 (n=38) and PND9 (n = 15-21). *, *p* < 0.05, **, *p* < 0.01, ***, *p* < 0.001.

Preterm pigs were delivered by cesarean section at 90% gestation from 8 sows (3 litters for 5-d study, 3 litters for 9-d study, 2 litters for 19-d study). All preterm pigs were block randomized according to the birth weight and sex and allocated into two groups by receiving either 0.5 mg/kg recombinant human IGF-1 with binding protein 3 (rhIGF-1/IGFBP-3) to serve as preterm IGF-1 group, or equivalent volumes of vehicle to serve as preterm control group. The rhIGF-1/IGFBP-3 supplementation for 5-d and 9-d studies was given by subcutaneous injections twice daily with the molar ratio of rhIGF-1/IGFBP-3 as 1:1 as described in our previous studies ^32,33^. For the 19-d study, the rhIGF-1/IGFBP-3 supplementation was administrated intraarterially through umbilical artery catheters during the first 7 days and switched to subcutaneous injections three times daily beyond postnatal day 7. Pigs were immunized and nourished with parenteral nutrition and infant formula as described in our previous studies ^33^. Vaginally born term reference pigs (Landrace x Yorkshire x Duroc, 117 ± 2 days) were raised at the farm by the sow until euthanasia. Term reference pigs were euthanized by matching the postnatal ages with preterm groups on postnatal days (PND) PND5, PND9 or PND19. Kidney tissue, blood and urine from all three groups were collected at euthanasia. Briefly, the kidneys were cut longitudinally from the middle. Half of the tissue was fixed in phosphate buffer paraformaldehyde for further histological staining, while another half was snap-frozen with liquid nitrogen and stored at -80□ for further analysis. After anesthesia, blood was collected with a heparin vacutainer (BD Diagnostics, Oxford, UK) via heart puncture sampling and urine was obtained through cystopuncture. All available kidney, blood and urine samples were included for related analysis. Information concerning IGF-1 contents in the circulation is shown in Supplemental Table S2. IGF-1 pharmacokinetics in piglets was reported previously ^32,33^.

During euthanasia, macroscopic indicators of Necrotizing Enterocolitis (NEC) were assessed by two independent observers, based on the presence of symptoms in the stomach, proximal, middle, and distal small intestine, and colon as previously described ^32,34^. Representative images of NEC scoring evaluation are displayed in Figure 1B. In order to further examine the effect of NEC on the kidney, preterm pigs on PND 5 were subdivided into three groups according to the NEC severity: the No NEC group (all score = 1), the mild NEC group (highest score in all gut region = 2-3), and the severe NEC group (highest score = 4-6). Furthermore, all preterm pigs in 5-d study were subgrouped according to lesion types into 3 groups: No lesion (all scores ≤2), small intestinal (SI) lesion (lesion score from SI ≥3), and non-SI tract lesion (lesion score from stomach and colon ≥3, and lesion score from SI ≤2). Kidney parameters were compared among groups to investigate the influence of NEC severities and lesion types on renal health.

### Biochemical analysis

Plasma and urine biochemistry was analyzed using an Advia 2120i Hematology system (Siemens Healthcare, Ballerup, Denmark). Conventional markers for kidney function assessment were included in the biochemistry profiling.

### Tissue Histology

Formalin fixed kidney tissues were embedded in paraffin, sectioned at 4 μm, then stained by hematoxylin and eosin (H&E) and periodic acid–schiff stain (PAS). H&E sections were used to measure and calculate the nephrogenic zone width (NZW) according to ^35^. PAS sections were used to measure mean glomerular area ^36^ and cross-sectional density ^37-39^, which reflect relative glomerular size and density in kidney tissues, respectively. Briefly, 50 complete glomeruli with clear capillary tuft and Bowman space were randomly selected for each section to measure and calculate the mean glomerular area. For cross-sectional density, 3 micrographs containing the nephrogenic zone and cortex area from each section were randomly taken under 50-fold magnification. Then all complete glomeruli were counted in micrographs to calculate the density. Meanwhile, abnormal glomeruli with shrunken tufts were counted to calculate the abnormal glomeruli percentage ^10,40^. Since injured vasculature in kidney is proven closely related to the deposition of extracellular matrix ^41^, the fractional mesangial area (FMA) was adopted to quantify the percentage of PAS-positive mesangial matrix within the glomerular tuft to reflect injury in the glomeruli. The FMA was measured under 400-fold magnification according to ^42^. All histological analyses were done by imageJ software version 1.50i (NIH, Bethesda, MD).

### Gene expressions by RT-qPCR

Transcription levels of genes related to kidney development, injury and inflammation were determined by RT-qPCR. Primers applied are shown in Supplemental Table S1. Briefly, total RNA was extracted from homogenized kidney cortex by using RNeasy Mini Kit (Qiagen, Copenhagen, Denmark). RNA was converted to cDNA using High-Capacity cDNA Reverse Transcription Kit (Thermo Fisher Scientific, United States). RT-qPCR was then performed with QuantiTect SYBR Green PCR Kit (Qiagen) on a LightCycler 480 (Roche, Hvidovre, Denmark), and relative expression of target genes was normalized to the housekeeping gene HPRT1 ^43^.

### Enzyme linked immunosorbent assay

Frozen kidney cortex was homogenized in cold lysis buffer and supernatant was obtained after centrifugation. Then the tumor necrosis factor alpha (TNFA) and interleukin 10 (IL10) protein levels in the tissue were measured by enzyme linked immunosorbent assay (ELISA) (R&D Systems, Abingdon, UK).

### Data analysis and statistics

Using R studio 3.6.1 (R Studio, Boston, MA, United States), univariate analysis was applied to all RT-qPCR, ELISA, biochemical and histologic quantification data at different time points of sampling. Each parameter was fitted to a linear mixed effect model by lme4 package with group and sex as the fixed factors, and litter as the random factor. Data were transformed when original data did not fit properly into the model by checking the residuals. If data could not be fitted to the model properly, then non-parametric analysis (Kruskal-Wallis test in package rstatix) was applied. Binary data were analyzed by a logistic regression model. NEC scores, as an ordered categorical outcome, were analyzed by a proportional odds logistic regression model. A *p*-value < 0.05 was regarded as statistically significant. Data are presented as mean ± SEM.

## Result

### Clinical outcomes

An overview of clinical parameters is displayed in Table 1. Term pigs had significant higher body weight and kidney weight at euthanasia compared to preterm pigs from both group (*p* < 0.001). The bodyweight and kidney weight at euthanasia were similar between preterm control and preterm IGF-1 groups. The incidence of severe NEC and distribution of digestive tract lesion in preterm pigs are displayed in Figure 1B-D. The NEC scoring criteria was described in our previous publication ^33^. The preterm control group had a higher incidence of severe NEC compared to the preterm IGF-1 group on PND5 (*p* < 0.05) and PND9 (*p* > 0.05). On PND5, the lesion score of the stomach and colon in the preterm control group was significantly higher than in the preterm IGF-1 group (both *p* < 0.05). The impact of varying severities or types of NEC lesions, with or without small intestine (SI) involvement has little effects on kidney health related parameters (Supplemental Figure S1 and S2).

**Table 1.** Body and kidney weight at birth and euthanasia in preterm control, preterm IGF-1 and term pigs.

|  | Group |  |  | *p*-value |  |
| --- | --- | --- | --- | --- | --- |
|  | Preterm control | Preterm IGF-1 | Term | Preterm control/Preterm IGF-1 | Preterm control/Term |
| Day 5 | n=37 | n=24 | n=7 |  |  |
| Birth weight (g) | 1067.7±51.6 | 1164.4±61.6 | - | 0.893 | - |
| Euthanasia weight (g) | 1136.8±53.9 | 1232.3±64.6 | 1688.1±69.9 | 0.999 | <0.001 |
| Kidney weight (g) | 10.1±0.5 | 10.8±0.7 | 13.7±0.8 | 0.565 | 0.006 |
| Day 9 | n=15 | n=21 | n=7 |  |  |
| Birth weight (g) | 947.9±74.1 | 953.8±54.4 | - | 0.802 | - |
| Euthanasia weight (g) | 1238.8±86.0 | 1269.8±84.4 | 3229.4±244.6 | 0.969 | <0.001 |
| Kidney weight (g) | 11.9±1.0 | 13.5±1.2 | 27.3±2.2 | 0.676 | <0.001 |
| Day 19 | n=13 | n=16 | n=7 |  |  |
| Birth weight (g) | 995.7±50.0 | 903.0±48.4 | - | 0.169 | - |
| Euthanasia weight (g) | 1976.6±70.3 | 1790.3±74.9 | 4491.4±457.6 | 0.600 | <0.001 |
| Kidney weight (g) | 14.0±0.8 | 14.5±0.8 | 33.0±3.1 | 0.931 | <0.001 |

### IGF-1 treatment improves kidney development following preterm birth

Overall, IGF-1 supplementation significantly increased the relative kidney weight of preterm pigs on PND19 (*p* < 0.05) (Figure 2A). The renal microstructure was examined by histological quantification to further investigate kidney maturation (Figure 2). The cross-sectional glomerular area in term pigs was significantly greater than in preterm control and IGF-1 groups at all time points (*p* < 0.01, Figure 2C). In contrast, the cross-sectional glomerular density was significantly lower in term pigs compared to preterm control and preterm IGF-1 groups on PND19 (*p* < 0.001, Figure 2D). The percentage of abnormal glomeruli, characterized by shrunk glomerular tufts on the outer renal cortex (Figure 2E) was significantly higher in the preterm control group on PND5 compared to the preterm IGF-1 treated group (*p* < 0.05) and term group (*p* < 0.001) (Figure 2F).

**Figure 2.**
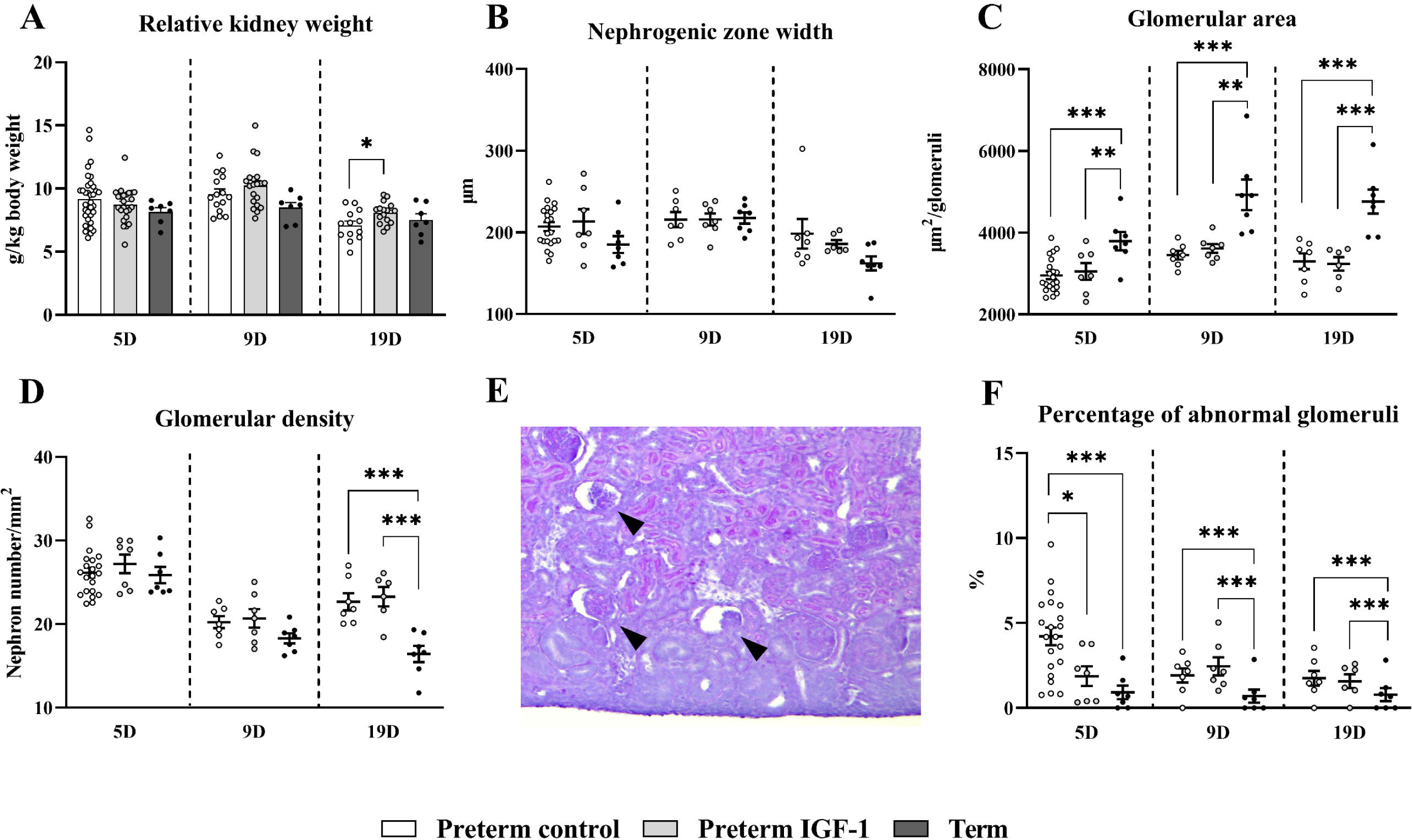
Effects of preterm birth associated immaturity and IGF-1 supplementation on kidney structure and maturation during the first 19 postnatal days. (A) Kidney weight relative to the bodyweight. (B) Nephrogenic zone width, measured with kidney H&E staining. (C) Cross-sectional glomerular area. (D) Glomerular density quantified with kidney PAS staining. (E) The representative picture from preterm control group indicates abnormal glomeruli (as cystic dilation of the Bowman’s space and shrunken glomerular tuft) appearing in the outer renal cortex. (F) The percentage of abnormal glomeruli. All data in preterm control (n=7-36), preterm IGF-1 (n=7-24) and term reference pigs (n=6-7) is presented as means ± SEM. *, *p* < 0.05, **, *p* < 0.01, ***, *p* < 0.001.

The expression of renal development-related genes was evaluated to assess the impact of preterm birth and IGF-1supplymentaion on kidney maturation. Preterm pigs showed a potential altered kidney development pattern as evidenced by the distinct expression pattern of SIX homeobox 2 (*SIX2*), *VEGFA*, *GDNF*, transforming growth factor beta 1 (*TGFB1*), Wnt family member 4 (*WNT4*), *WNT9B*, *WNT11*, and E-cadherin (*CDH1*) compared to term pigs (Figure 3). Furthermore, IGF-1 supplementation significantly upregulated the expression of *GDNF* (*p* < 0.001), *TGFB1* (*p* < 0.01), angiotensin II type 1 receptor (*AT1, p* < 0.05) and catenin beta 1 (*CTNNB1, p* < 0.05) on PND5 compared to the preterm control. However, no differences of above expression were detected on PND9 and PND19 between preterm IGF-1 and preterm control groups. These findings provide further evidence of the role of IGF-1 supplementation in early-stage kidney development after preterm birth.

**Figure 3.**
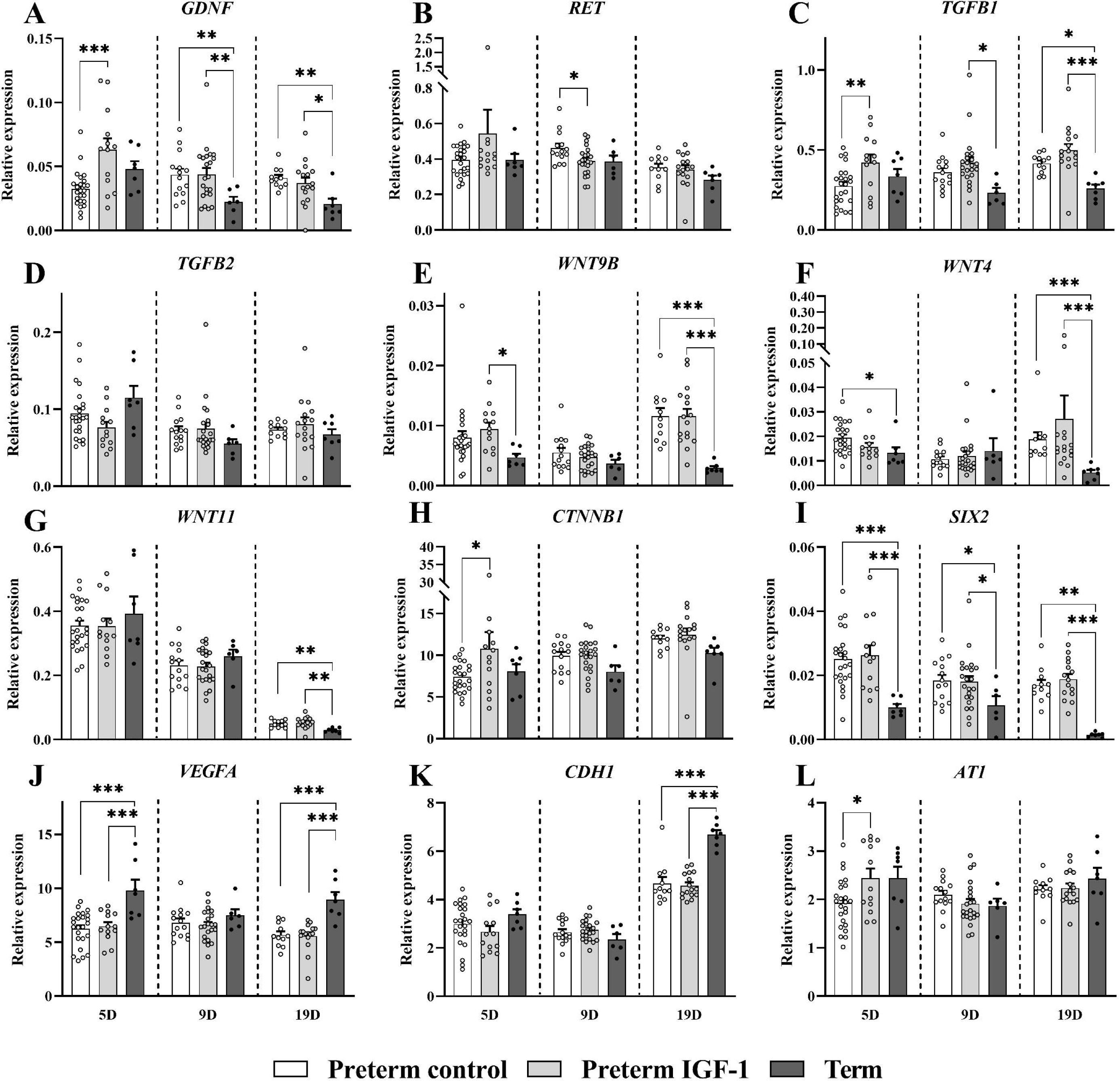
Effects of preterm birth associated immaturity and IGF-1 supplementation on development related gene expression during the first 19 postnatal days. (A and B) Glial-cell-line-derived neurotrophic factor/Ret proto-oncogene signaling molecules (*GDNF* and *RET*), (C and D) transforming growth factor-beta (*TGFB1* and *TGFB2*), (E-H) Wnt family members and their downstream molecule β-catenin *(WNT4*, *WNT9B*, *WNT11* and *CTNNB1*), (I) nephron progenitor marker *SIX2*, (J-K) other kidney development related genes (*VEGFA*, *CDH1*, and *AT1*) in kidney tissues. All data in preterm control (n=11-24), preterm IGF-1 (n=12-22) and term reference pigs (n=6-7) is presented as means ± SEM. *, *p* < 0.05, **, *p* < 0.01, ***, *p* < 0.001.

### IGF-1 treatment may ameliorate the kidney injury associated with preterm birth

The kidney function and injury status were analyzed by examining plasma and urine biochemistry as shown in Figure 4. On PND5, preterm pigs exhibited a higher level of plasma creatinine than term pigs, after which the trend reversed. Moreover, blood urea nitrogen (BUN) level in term group was higher than the two preterm groups on PND5 (*p* < 0.001) and PND19 (*p* < 0.05). The plasma albumin level was consistently higher in term pigs than preterm pigs at all time points (*p* < 0.05). The biochemistry parameters tested were similar between preterm control and IGF-1 pigs.

**Figure 4.**
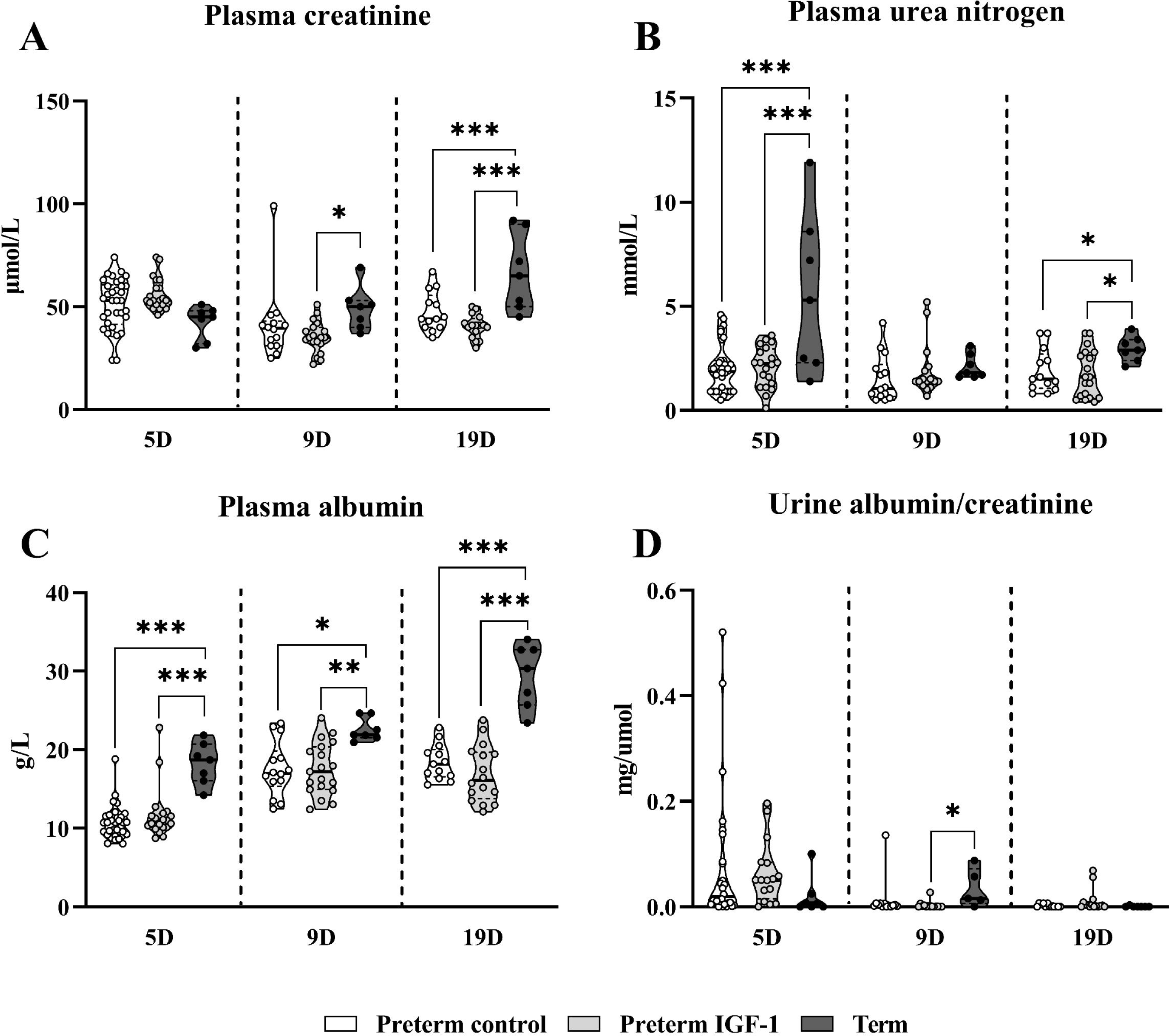
Effects of preterm birth associated immaturity and IGF-1 supplementation on plasma and urine biochemical parameters during the first 19 postnatal days. (A) Plasma creatinine level; (B) Plasma urea nitrogen level; (C) Plasma albumin level; (D) Urine albumin to creatinine ratio. All data in preterm control (n=12-34), preterm IGF-1 (n=14-20) and term reference pigs (n=6-7) is presented as means ± SEM. *, *p* < 0.05, **, *p* < 0.01, ***, *p* < 0.001.

To determine if there was any potential kidney injury, the study measured the expression of kidney injury-related genes, as shown in Figure 5. The expression of kidney injury molecule-1 (*KIM1, p* < 0.05) and leucine-rich alpha-2-glycoprotein 1 (*LRG1, p* < 0.01) was significantly higher in the preterm control group than in preterm IGF-1 and term pigs on PND5 (Figure 5A and 5B), suggesting a potential protective effects of IGF-1 supplementation on preterm birth related kidney injury. The FMA of the glomeruli, which indicates glomeruli injury, was consistently higher in the preterm control and IGF-1 groups than the term group at all time points (*p* < 0.05). The IGF-1 treated group showed slightly lower FMA levels compared to vehicle treated preterm pigs on PND5 (*p* > 0.05, Figure 5C and 5D).

**Figure 5.**
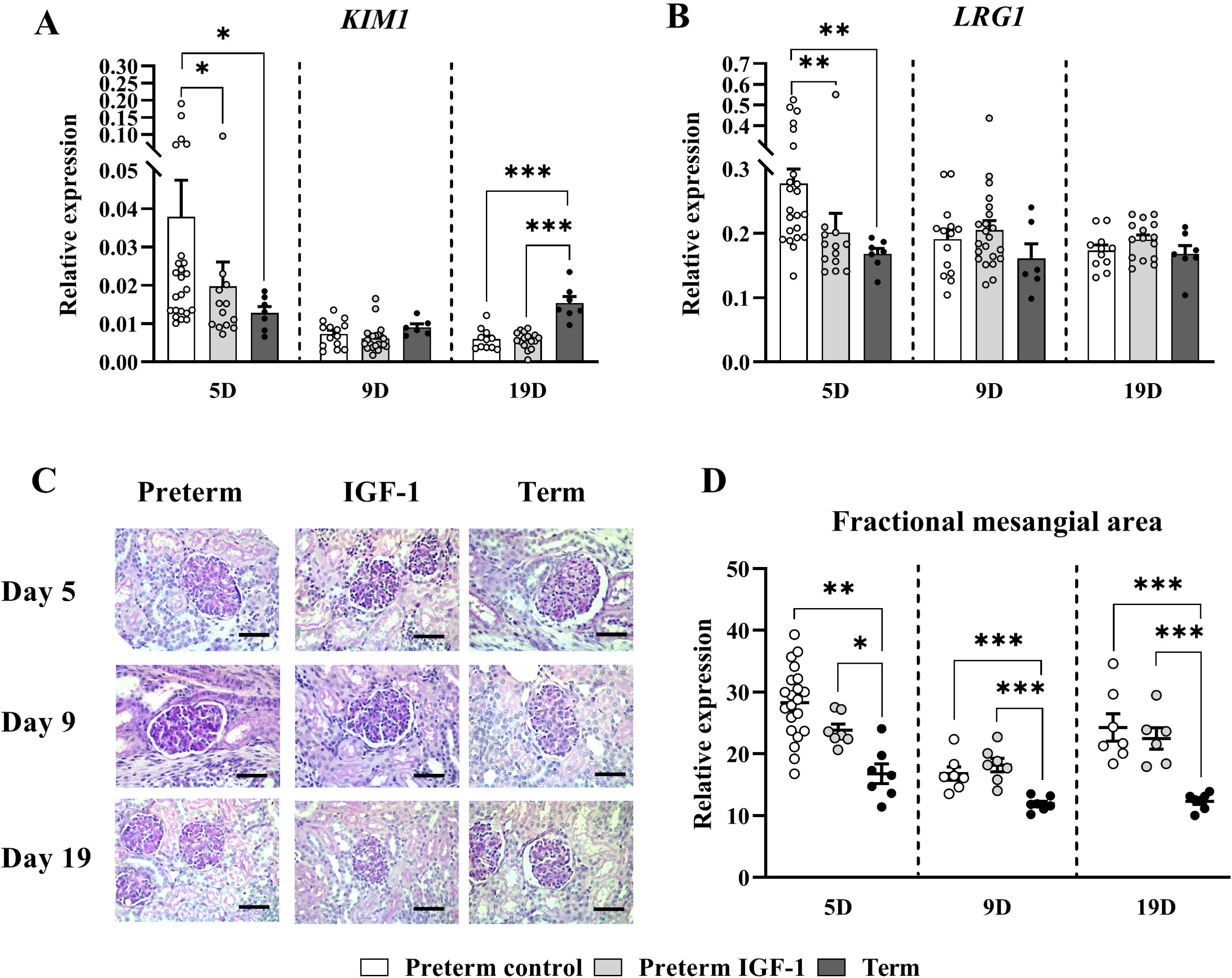
Effects of preterm birth associated immaturity and IGF-1 supplementation on kidney injury related parameters during the first 19 postnatal days. (A and B) Relative gene expression of kidney injury markers (KIM1 and LRG1) in renal tissue. (C) Representative histological images of the glomeruli in preterm, IGF-1 and term groups at three time points (PAS staining, scale bars: 50 μm). (D) Fractional mesangial area of the glomeruli at three time points. All data in preterm control (n=7-21), preterm IGF-1 (n=7-22) and term reference pigs (n=6-7) is presented as means ± SEM. *, *p* < 0.05, **, *p* < 0.01, ***, *p* < 0.001.

### IGF-1 treatment ameliorated preterm birth associated kidney inflammation

As shown in Figure 6, preterm birth was associated with elevated inflammatory responses in the kidney on PND 5, as indicated by dramatically upregulated renal gene expression of *TNFA* and *IL10* in the preterm control group, relative to the term group (*p* < 0.05 and *p* < 0.001, respectively). Treatment with IGF-1 resulted in a significant downregulation in *TNFA* (*p* < 0.05) and *IL10* (*p* < 0.001) expression in the kidney compared to the preterm control group on PND 5 (Figure 6A and 6B). Moreover, the renal protein levels of TNFA and IL10 exhibited a similar trend to their mRNA levels among groups on PND 5 (Figure 6D and 6E). *IL6* gene expression showed no significant differences among all groups (*p* > 0.05, Figure 6C).

**Figure 6.**
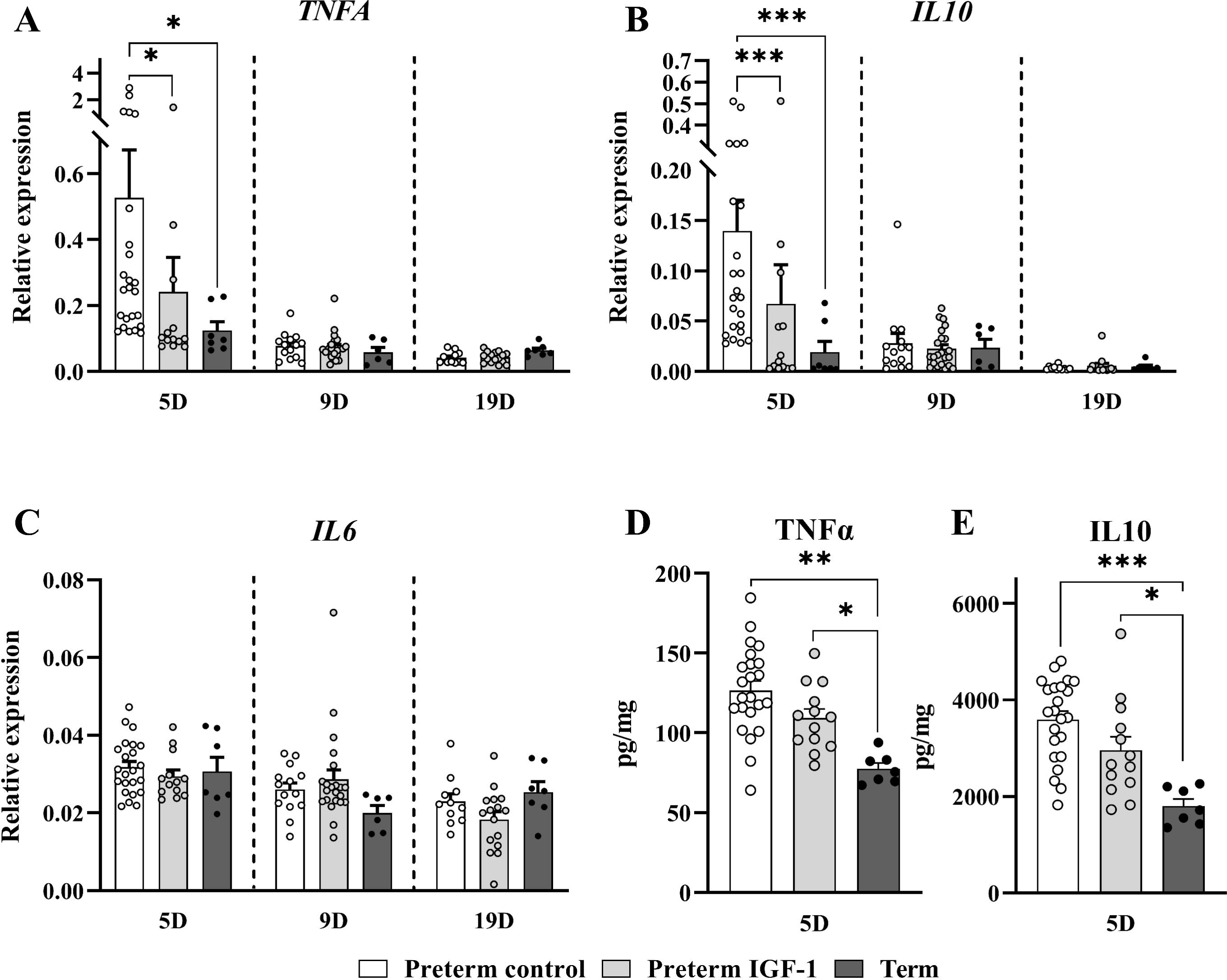
Effects of preterm birth associated immaturity and IGF-1 supplementation on inflammation related parameters during the first 19 postnatal days. (A, B and C) Relative gene expression of three inflammatory cytokines (TNFA, IL10 and IL6) in kidney tissue. (D and E) Renal protein expression of two inflammatory cytokines (TNFA and IL10). All data in preterm control (n=11-24), preterm IGF-1 (n=12-22) and term reference pigs (n=6-7) is presented as means ± SEM. *, *p* < 0.05, **, *p* < 0.01, ***, *p* < 0.001.

Pearson correlation analyses were used to explore the potential association between kidney inflammatory response and injury status. The analysis revealed a significant positive correlation between the expression of genes associated with kidney inflammation and those related to kidney injury at PND5 (Supplemental Figure S3), suggesting that preterm birth-associated kidney injury may be associated with elevated local inflammatory status.

## Discussion

In preterm infants, low levels of circulating IGF-1 are associated with fetal and postnatal growth restriction, systemic inflammation and multiple organ-related complications ^33^. A large multicenter international trial presently is underway to determine whether the administration of supplemental IGF-1 can reduce morbidities in extremely preterm infants (ClinicalTrials.gov registry NCT03253263), partly based on the initial evidence of reduced bronchopulmonary dysplasia and intraventricular hemorrhage ^44^. In this study, preterm pigs were utilized as a model to further assess the clinical safety and efficacy of rhIGF-1/IGFBP-3, targeting a series of organ systems. Prior investigations from our group revealed that supplementary IGF-1 had a moderate impact on gut development and NEC preventing after a treatment period of 5-9 days ^32,33^. In the current study, we show that supplemental IGF-1 treatment may promote nephrogenesis and alleviate preterm birth associated renal inflammation and injury. Correlation analyses between kidney variables and NEC lesion scores suggest that the effects of IGF-1 treatment on immature kidney do not interact with gut-related inflammation, at least not when NEC lesions are relatively mild without systemic effects. We conclude that systemic IGF-1 supplementation exerts direct effects on renal health in preterm newborns.

Nephrogenic zone is an active region in the outer layer of the cortex where immature and primitive glomeruli develop, the width of which shows a negative correlation with gestational age at birth ^9,35^. Moreover, studies in humans suggest nephron density is negatively correlated with kidney maturation stage probably due to higher volume of interstitial tissue ^38^. The increase of NZW, and the decrease of glomerular area ^39^ and density all indicate a delayed postnatal kidney maturation in preterm piglets compared to term pigs. Further, the differential expression pattern of renal development associated genes, including elevated levels of *GDNF*, *Wnt* family members, and *SIX2* (a nephron progenitor marker) on PND19 in preterm pigs, might suggest continued immaturity and maturation activity of preterm kidney ^45^.

Preterm newborns are more susceptible to AKI during the perinatal period, typically due to immature renal structure and function ^12,46-48^. Clinically, AKI commonly occurs within the first week of postnatal life in preterm newborns ^49-51^. This corresponds to our finding that the kidney of preterm pigs is most vulnerable and sensitive to insults during the first 5 days of postnatal life. The preterm newborn kidney may recover over time partly due to adaptation to the new physiological and environmental conditions. Intra-renal inflammatory responses are present in AKI conditions ^52^. Ischemia-reperfusion injury, a leading cause of AKI, induces inflammation in renal endothelial cells ^53^.

The causes of AKI in preterm infants are multifactorial and can arise from systemic conditions ^54^. NEC is a rapidly escalating gastrointestinal inflammatory disease that occurs in preterm infants in the first few weeks after birth ^20,21,55^. Severe NEC lesions, also known as surgical NEC, may predispose preterm infants to AKI ^23,56^, although the underlying mechanisms remain unclear. Critically ill preterm neonates with severe NEC symptoms may experience AKI due to their critical condition, sepsis and hemodynamic instability. While if NEC lesions are an independent nephrotoxic factor is unknown ^57^. A mild idiopathic NEC animal model is hired in this study to investigate the potential association. In this preterm pig model, the predisposing factor for NEC was preterm birth combined with formula feeding, consistent with the key risk factors leading to NEC in infants ^58^. As we found limited evidence of correlations between kidney insults and NEC lesions (here clinically defined as mild NEC in our model), we conclude that only severe NEC lesions, which require surgical intervention for survival in both pigs and infants ^23,59^, may predispose preterm neonates to kidney injury in the first weeks of life. Although a study in mice showed hypoxia-induced NEC led to AKI, the systemic hypoxia can directly cause renal damage ^60^. Recent clinical studies have reported that kidney failure only occurred in 2.8% of infants with NEC ^61^. Similar low incidences of NEC (< 2%) have also been observed in clinical studies on AKI ^62^, including the large Acute Kidney Injury Epidemiology in Neonates study ^57^.

Studies have demonstrated that IGF-1 promote kidney growth and development during the neonatal period ^31,63^, which potentially explain the enlarged kidney observed in preterm IGF-1 pigs on PND19. Morphologically abnormal glomeruli, characterized by cystic dilation of the Bowman’s space and shrunken glomerular tufts in the superficial renal cortex, were documented in preterm baboons, lambs as well as infants ^9,10,64,65^. These glomeruli are the last to be formed and believed to result from an accelerated adaptation to the extrauterine environment. In this study, the increased percentage of abnormal glomeruli in preterm versus term pigs confirms the impact of shortened gestational length on postnatal kidney development. While the administration of IGF-1 appears to support the morphological structure of the last formed glomeruli. Both the GDNF/RET ^16,66^ and Wnt/β-catenin ^67,68^ pathways are essential in regulating mesenchymal-epithelial transformation and branching morphogenesis in the kidney. TGFβ1 is involved in elongation of the ureteric bud and nephron form ^69,70^, while *AT1* is associated with encoding various components of cytoskeleton and extracellular matrix during normal neonatal renal development ^71^. The upregulation of *GDNF*, *RET*, *CTNNB1*, *TGFB1* and *AT1* expression in the kidney after IGF-1 treatment on PND5 suggests that IGF-1 may benefit or accelerate essential processes of nephrogenesis in preterm pigs immediately after birth. Besides, AT1, as a component of the renin–angiotensin system, plays a crucial role in regulating blood pressure and renal hemodynamics ^72^. Hence the regulation of *AT1* by IGF-1 may support maintaining stable hemodynamics, which warrants further investigation ^29,38^. In addition, IGF-1 may accelerate recovery from kidney injury and protect against acute failure, probably via its anti-apoptotic and mitogenic properties ^73-76^. In the current study, such protective effects were most pronounced during the first week of life. Given that neonatal AKI may lead to reduced glomerular filtration rate (GFR), glomerulosclerosis and chronic kidney disease later in life ^77^, early treatment of preterm infants to prevent urther negative renal impacts is critical.

While these findings provide new insights into neonatal kidney injuries associated with preterm birth, our analysis of this study has certain limitations. The primary limitations of the present study include the lack of confirmation of kidney injury and dysfunction through direct or conventional evidence such as decreased GFR, changed plasma creatinine coupled with urine output. However, it should be noted that plasma creatinine levels during the neonatal period are not always reliable for defining AKI as they primarily reflect maternal creatinine levels ^78,79^. Although increased BUN levels during early life are considered as an indicator for immature renal function ^80^, BUN levels are primarily affected by protein intake and store, and amino acid metabolic levels, which may obscure alterations in renal function ^81^. In the present study, elevated BUN levels in term pigs may be partly due to their higher protein intake and store under natural suckling conditions. Moreover, the effects of IGF-1 require further validation as several variables showed limited differences between preterm control and IGF-1 groups. Hence, further investigations to assess the short- and long-term physiological and functional outcomes of IGF-1 treatment are warranted (e.g. GFR, blood pressure, fluid balance). Despite these limitations, we conclude that immediate postnatal IGF-1 supplementation may promote renal development and alleviate renal inflammation and injury in preterm newborns. Possibly, supplemental IGF-1 could be most effective in situations just after preterm birth with high risk of systemic inflammation induced by prenatal infection/inflammation or severe postnatal inflammatory morbidities such as surgical NEC or late-onset sepsis.

## Data availability statement

All datasets generated during and analysed during the current study, including raw data used for all figures, tables and analysis, are available from the corresponding author on reasonable request.

## Author contributions

Conception and design: JZ, PTS, DNN, TM; data acquisition: JZ, TM; data analysis: JZ, TT, PJ, DNN, TM; data interpretation: JZ, PTS, DNN, TM; writing original draft: JZ, TM; critical review and editing: TT, PTS, DNN, PJ, TM; approval of the final manuscript: all co-authors.

## Conflict of interest

Author PTS, TT, DNN, and TM are currently involved in a patent application directed to use of rhIGF-1 for preterm infants. The remaining authors declare that the research was conducted in the absence of any commercial or financial relationships that could be construed as a potential conflict of interest.

## Funding

The authors declare that this study received funding from Takeda, MA, USA (grant number: B11035M-TAK607; grant recipient: Per Torp Sangild). The funder was not involved in the study design, execution, collection, analysis, interpretation of data, the drafting of this article or the decision to submit it for publication.

