## Supplement tables and figures for "Insulin-like Growth Factot-1 Supplementation Promotes Kidney Development and Alleviate Renal Inflammation in Preterm Pigs"

**Supplemental Table S1 List of gene names, primer sequences and amplicon length of primers used in the study.**

| **Gene name** | **Gene symbol** | **Sequence (5' to 3')** | **Sequence (5' to 3')** | **Amplicon length** |
| --- | --- | --- | --- | --- |
| HPRT1 | Hypoxanthine phosphoribosyltransferase 1 | TATGGACAGGACTGAACGGC | ATCCAGCAGGTCAGCAAAGA | 115 |
| CASPASE3 | Caspase 3 | CCGGAATGGCATGTCGATCT | CATGGCTTAGAAGCACGCAA | 189 |
| GATA3 | GATA binding protein 3 | ACCCCTTATTAAGCCCAAGC | TCCAGAGAGTCGTCGTTGTG | 92 |
| IL10 | Interleukin 10 | GTCCGACTCAACGAAGAAGG | GCCAGGAAGATCAGGCAATA | 73 |
| IL6 | Interleukin 6 | TGGGTTCAATCAGGAGACCT | CAGCCTCGACATTTCCCTTA | 116 |
| TBET | T-box transcription factor 21 | CTGAGAGTCGCGCTCAACAA | ACCCGGCCACAGTAAATGAC | 121 |
| TNFA | Tumor necrosis factor alpha | ATTCAGGGATGTGTGGCCTG | CCAGATGTCCCAGGTTGCAT | 120 |
| HIF1A | Hypoxia-inducible factor 1 alpha | TGTGTTATCTGTCGCTTTGAGTC | TTTCGCTTTCTCTGAGCATTC | 96 |
| KIM1 | Kidney injury molecule-1 | ATGTACCCTTGGGTAACCGC | AACGTAGAACATGCCCCTCG | 164 |
| NGAL | Neutrophil gelatinase-associated lipocalin | AAGACGGCAGCTACAACGTC | GACACCACACGCACGACATA | 157 |
| LRG1 | Leucine Rich Alpha-2-Glycoprotein 1 | TGACCTGCACATCCTTGACC | CAGAAAGCCCTCTTCGAGCA | 133 |
| IGFBP7 | Insulin like growth factor binding protein 7 | ATCGTGACACCCCCTAAGGAC | ATAGTGACCCCTTTGTACCTTGT | 120 |
| WNT11 | Wnt family member 11 | CTTGACCTGGAGAGAGGGACC | ATGAGGAGCCCGTAGCTGAG | 197 |
| WNT4 | Wnt family member 4 | CACAAGGCTTCCAGTGGTCA | GCATGTGTGTCAGGATAGCCTTC | 168 |
| WNT9b | Wnt family member 9B | CTGCTCGAGTGCCAGTTTCA | CCGTCTCCTTGAAACCTCTCTT | 94 |
| VEGFA | Vascular endothelial growth factor A | ATGCGGATCAAACCTCACCA | TGTCACATCTGCAAGTACGTTCG | 224 |
| TGFB1 | Transforming growth factor beta 1 | GCAAGGTCCTGGCTCTGTA | TAGTACACGATGGGCAGTGG | 97 |
| TGFB2 | Transforming growth factor beta 2 | GCGCGATTTGCAGACTTGAG | ATGTAAAGTGGACGCAGGCA | 170 |
| REN | Renin | TCTCCGTCTACTACAGCAGGA | CCTCACAGACACCCCTTTCA | 152 |
| RET | Ret proto-oncogene | GGAAGATAACCAGGACCCGC | GTTCTTGTGGTAGCGGTGGA | 124 |
| SIX2 | SIX homeobox 2 | CTACCCCTCACCCCGAGAGA | TTCTCGCTGTTCTCCTCGTACC | 141 |
| CTNNB1 | Catenin beta 1 | ATTGAAGCTGAGGGAGCCAC | GAACTGGTCAGCTCAACCGA | 152 |
| AT1 | Angiotensin II receptor type 1 (AGTR1) | GGTCTACATCCAGGTGCATT | GGGGCAGTCATCTTGGATTC | 116 |
| CDH1 | E-cadherin | GCTGGACCGGGAGAGTTTTC | TGAAGATGGGTGGGTTGTCG | 131 |
| GDNF | Glial cell derived neurotrophic factor | CCTGGGCGTGTGGATGTTTA | GCCATCTGTTTATCGGGGGA | 132 |
| IGF-1 | Insulin-like growth factor 1 | ATTTCTTGAAGGTAAAGATGCA | CAGCCCCACAGAGGGTCTCA | 117 |

**Supplemental Table S2 Circulating IGF-1 contents (ng/ml) in the plasma of pigs.**

|  |  |  | **Group** | | |  | ***p*-value** | |
| --- | --- | --- | --- | --- | --- | --- | --- | --- |
|  |  |  | **Preterm** | **IGF-1** | **Term** |  | **Preterm/IGF-1** | **Preterm/Term** |
| **Day 5** |  |  | n=37 | n=24 | n=7 |  |  |  |
| **IGF-1 contents** | |  | 10.64±0.64 | 50.19±3.96 | 16.13±2.94 | | <0.001 | 0.048 |
| **Day 9** |  |  | n=15 | n=21 | n=7 |  |  |  |
| **IGF-1 contents** | |  | 18.74±2.69 | 87.63±11.71 | 46.87±7.15 | | <0.001 | 0.125 |
| **Day 19** |  |  | n=13 | n=16 | n=7 |  |  |  |
| **IGF-1 contents** | |  | 42.94±2.53 | 139.32±9.75 | 45.45±8.55 | | <0.001 | 0.999 |

**Supplemental Figure S1 NEC severities effects on renal parameters on postnatal day 5.**  (A) Relative kidney weight; (B) Urine albumin/creatinine ratio; (C) Abnormal glomeruli percentage; (D) Fractional mesangial area; (E-I) Relative expression of kidney injury and inflammation related genes; (J and K) Kidney protein expression of TNFA and IL10. All data from pigs of no lesions (score 1, n=9-19), mild lesions (score 2-3, n=5-17) and severe lesions (scores 4-6, n=11-27) is presented as means ± SEM. *, *p* < 0.05, **, *p* < 0.01, ***, *p* < 0.001.

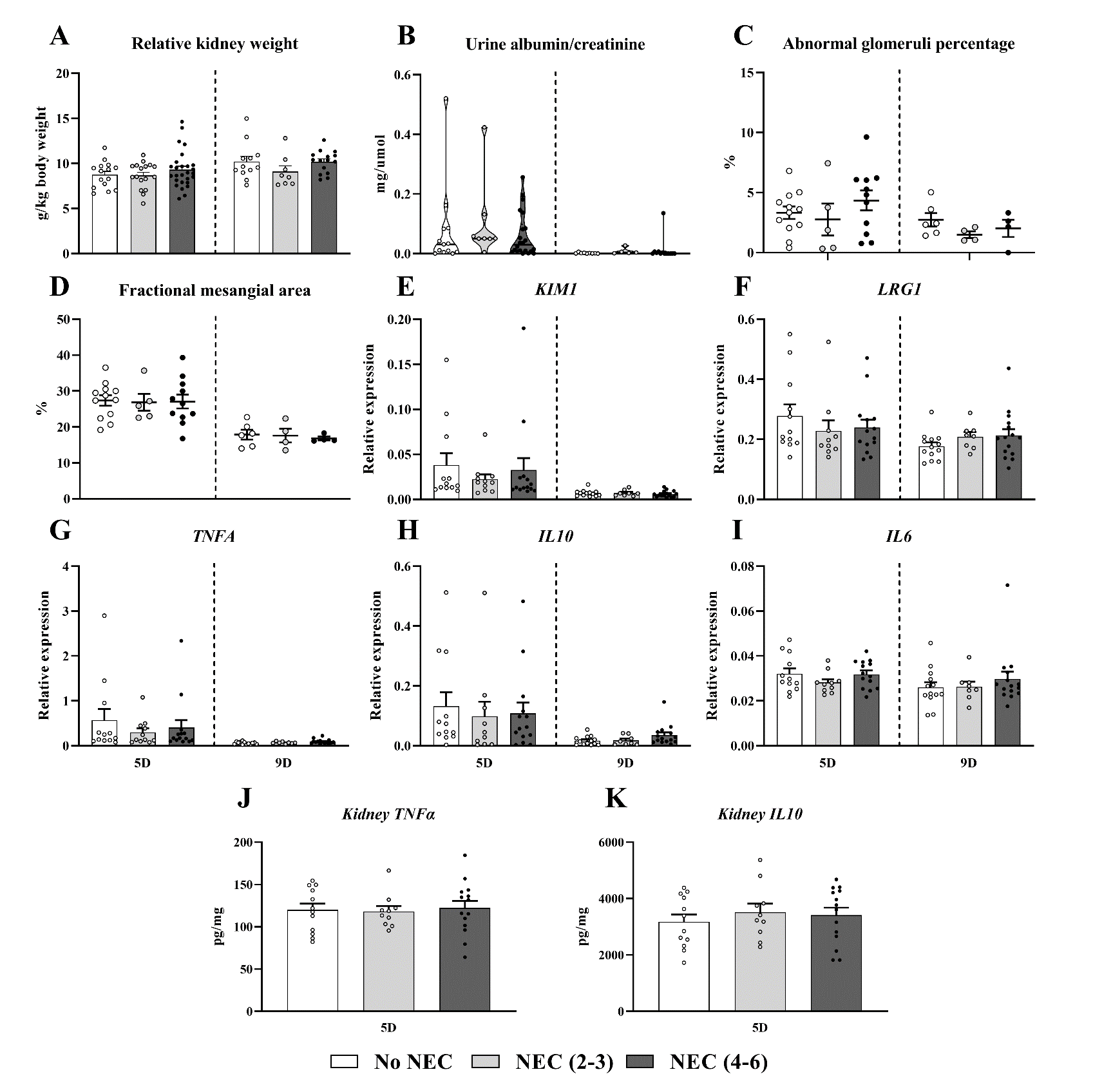

**Supplemental Figure S2 Effects of NEC lesion location type on renal parameters on postnatal day 5.** (A) Relative kidney weight; (B) Urine albumin/creatinine ratio; (C-G) Relative expression of kidney injury and inflammation related genes; (I and J) Kidney protein expression of TNFA and IL10. All data from pigs of no lesions (n=20-23), non-SI lesions (n=15-32) and SI lesions (n=5-7) is presented as means ± SEM. *, *p* < 0.05, **, *p* < 0.01, ***, *p* < 0.001.

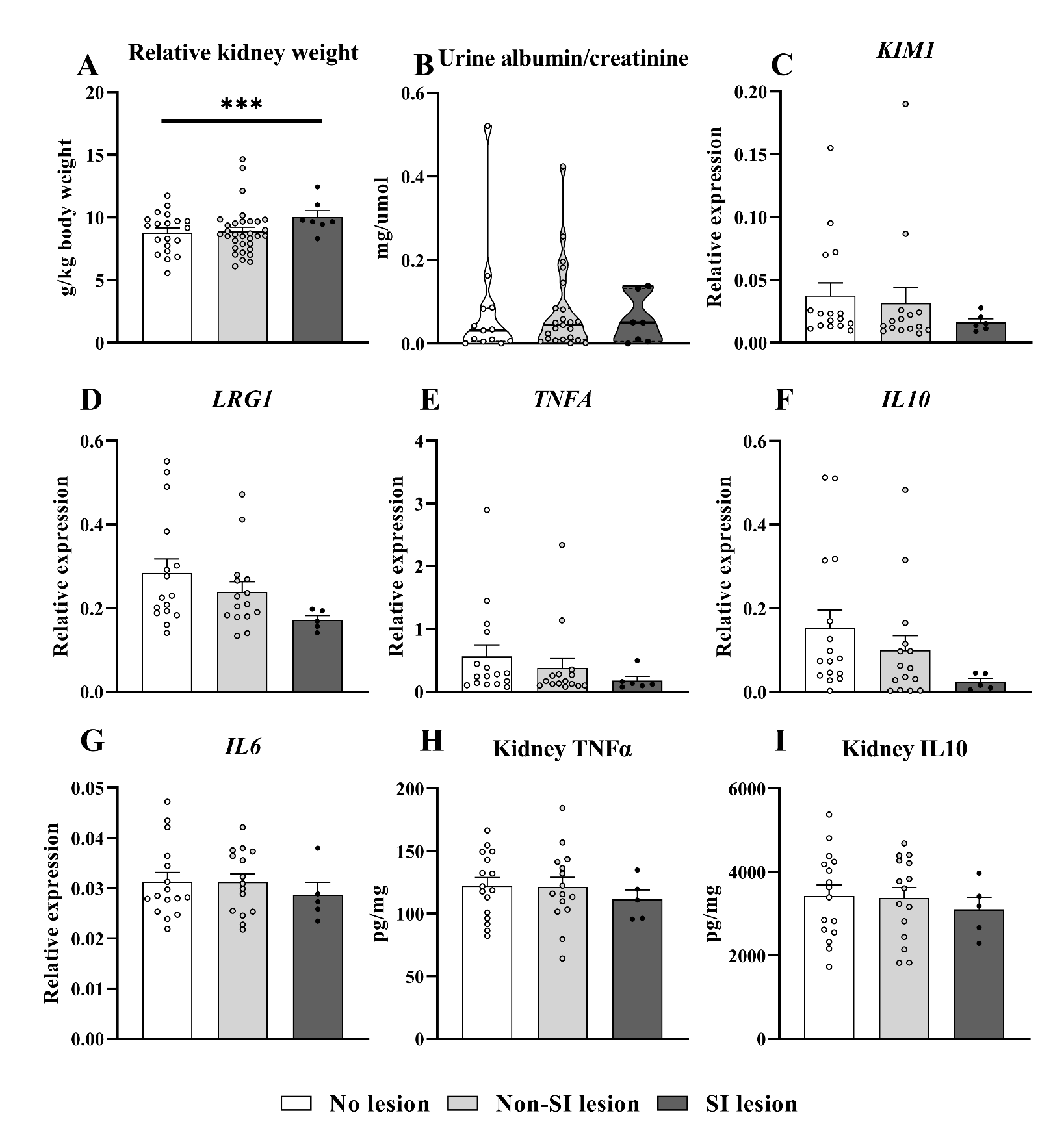

**Supplemental Figure S3 Pearson correlation analysis of kidney inflammation and injury related gene expression on PND5.** (A), (B), (C) and (D) are the correlation of *KIM1* to *TNFA*, *LRG1* to *TNFA*, *KIM1* to *IL10*, and *LRG1* to *IL10*, respectively.

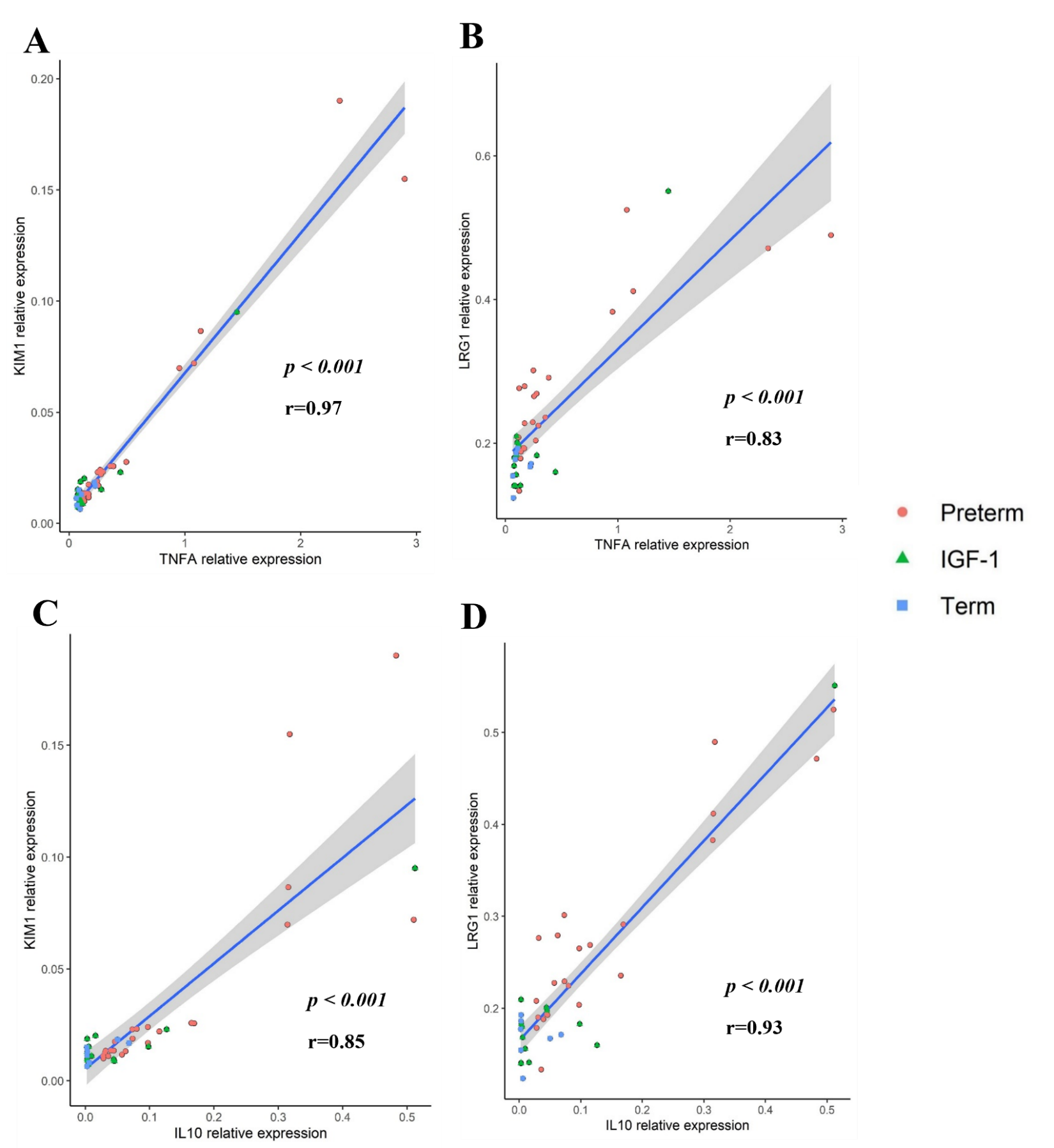
